# A simplified calculation of the correlations between relatives

**DOI:** 10.1101/2020.03.05.978361

**Authors:** Reginald D. Smith

**Affiliations:** Supreme Vinegar LLC

**Keywords:** correlation of relatives, quantitative genetics, kinship

## Abstract

The correlations between relatives is one of the fundamental ideas and earliest success of quantitative genetics. Whether using genomic data to infer relationships between individuals or estimating heritability from correlations of phenotypes amongst relatives, understanding the theoretical genetic correlations is a common task. Calculating the correlations between arbitrary relatives in an outbred population, however, can be a careful and somewhat complex task for increasingly distant relatives. This paper introduces an equation based method that consolidates the results of path analysis and uses easily obtainable data from non-inbred pedigrees to allow the rapid calculation of additive or dominance correlations between relatives even in more complicated situations such as cousins sharing more than two grandparents and inbreeding.

## 1 Introduction

One of the key achievements of the Modern Synthesis in evolutionary biology was the determination of the expected degree of genetic relationship between relatives given a known pedigree. First rigorously defined and expanded in Fisher’s landmark 1918 paper [Fisher 1918], the correlation of relatives became one of the first applied tools of quantitative genetics valuable for selective breeding and analysis of complex genetic disorders in families. Important contributions by Sewall Wright, especially path analysis [Wright 1922, Wright 1934] and Gustave Malécot [Malécot 1948, Malécot 1970] helped create the modern methods and notations to investigate the correlations between relatives for loci with additive or dominance contributions to genetic variance.

Further work on the correlation of relatives focused on the correlations due to epistatic effects [Kempthorne 1954, Cockerham 1954, Kempthorne 1955] including adjustments for linkage and linkage disequilibrium [Cockerham 1956, Gallais 1974]. While the correlations amongst relatives is well-known for almost all typical situations, the methods of calculating these correlations remain algorithmic and recursive. In this paper, an exact equation using the information from the structure of a known pedigree will be used to demonstrate a consistent mathematical formulations for the additive and dominance genetic correlations between relatives for all situations.

## 2 The relation of correlation and kinship to general pedigree variables

While several methods of calculating the correlation of relatives are known, almost all are recursive requiring the tracing of paths between relatives or calculating multiple types of identity coefficients in order to determine the relationship between two related individuals [Wright 1922, Gillois 1965, Jacquard 1972, Karigl 1981, Lange 2012]. However, once the correlations for various sets of relatives have been determined by analysis, it is possible to consolidate them into a general equation which combines previously recognized insights and understandings for different relative pairs.

Using notation similar to the original notation from [Wright 1922], for two relatives, *X* and *Y* with a set of most recent common ancestors *A*, let us define three variables: *G*_*XA*_, the number of generations of descent from any member of *A* to *X* and *G*_*Y A*_, the number of generations of descent from any member of *A* to *Y* . *G*_*XA*_ and *G*_*Y A*_ are *n* and *n*′ in [Wright 1922]. For half-sibs or half-cousins, one additional path is required through the parent of the halfsib/cousin so also define *H*(*X, Y*) as a binary variable that is one if either *X* or *Y* are half-siblings or half-cousins and zero otherwise.

Define as *C*, the number of elements of *A* which designates the number of common ancestors in the generation of *A*. For typical pedigrees *C* = 1 for direct descent (parents-offspring or grandparents-grandchildren) or *C* = 2 for other relatives. However, when cousins share more than two ancestors in *A*, such as double cousins sharing four grandparents, *C* can increase up to all ancestors for both relatives in *A* where the maximum value is reached at 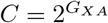. Finally, also including the possibility that the ancestral generation is inbred with coefficient *F*_*A*_, we can state the additive correlation between relatives *X* and *Y* in logarithmic form:

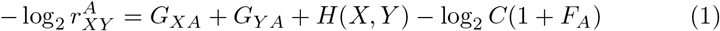

In the typical case where *C* = 2 and *F*_*A*_ = 0 this reduces to

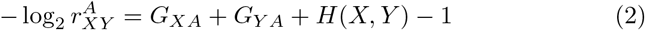

For cases where *X* and *Y* are inbred

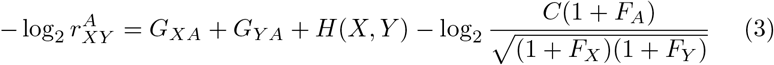

In Table 1, the actual and compared values of 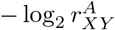 are shown for several situations where *C* = 1 or *C* = 2. Therefore using equation 1 one can quickly and accurately determine the correlation between relatives. The kinship coefficient is defined as 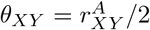.

**Table 1.**
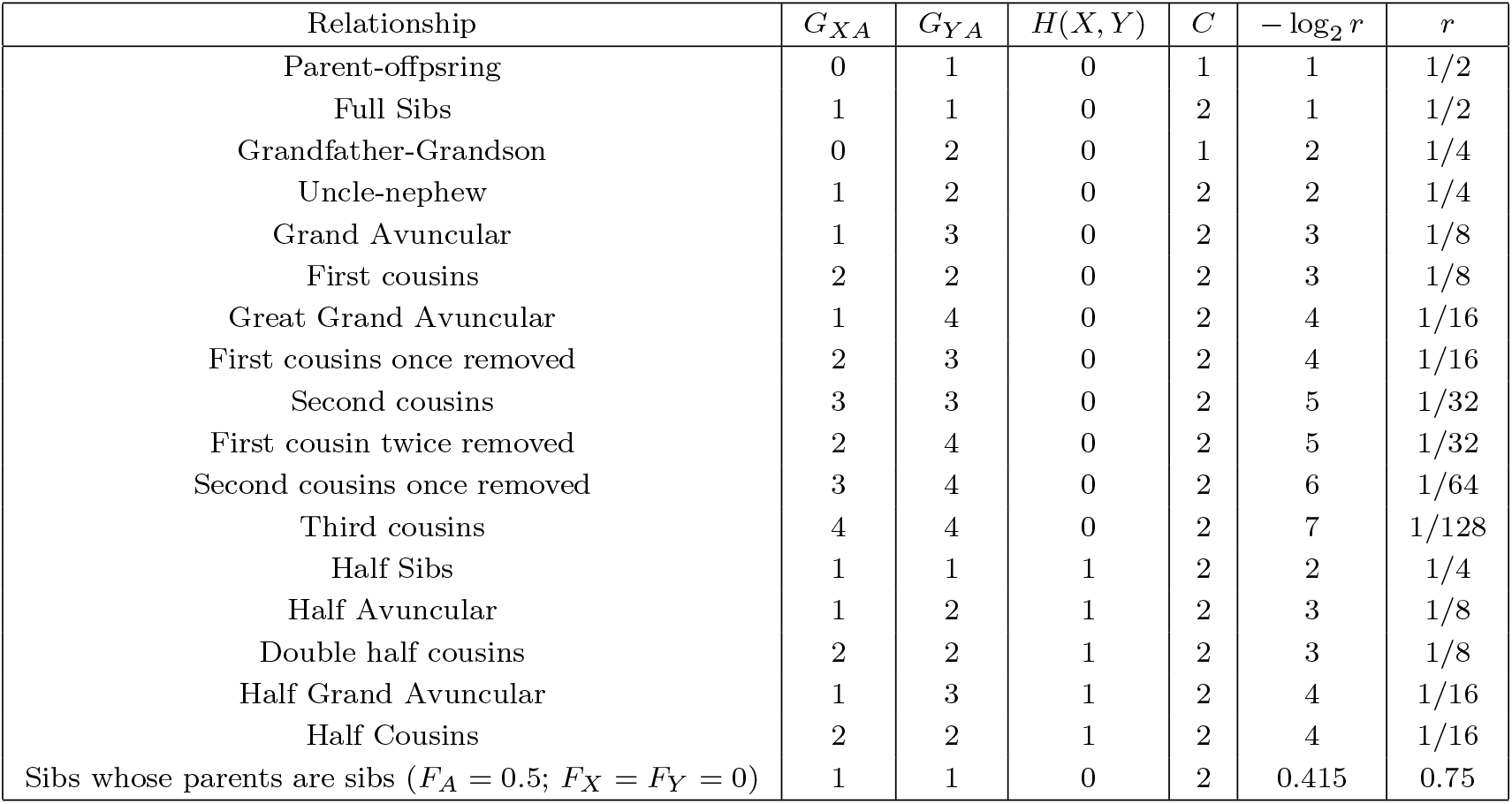
A basic table of relatives relating the variables derived from pedigrees and the additive correlations amongst the relatives. Values from [Karigl 1981] and [Ramstetter 2017].

### 2.1 Complex cousins

The more complicated situation arises when cousins have more than two relatives in common in the generation of their most recent common ancestor. For example, double first cousins share all four grandparents since each of their parents are from one of two pairs of full siblings. “Sesqui” first cousins share three grandparents due to one parent coming from a pair of full siblings and another coming from a pair of half siblings. Except double cousins, the terminology for such cousins is not universally standardized. For purpose of analysis, however, normal cousins of any degree should only share two relatives in the generation of their most recent common ancestors. Those that share more have an increased level of relationship that is reflected in their additive or dominance correlations. Using equation 1 we can derive the correlations for a variety of half or full cousins sharing more than one ancestor in the generation of *A*.

Having the ability to derive 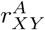, we can additionally calculate the dominance covariance and correlation in tandem with certain insights about the probability of sharing pairs of alleles that are identical by descent. Malécot used the insights of Fisher and Wright to derive the following expression of the total covariance between two relatives

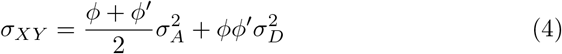

In equation 4, 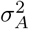 and 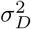 are the additive and dominant variance respectively, and *ϕ* and *ϕ′* are the correlations between each pair of loci in two relatives as to whether they both are represented by an allele that is identical by descent. For cousins, *ϕ* can be determined based on the number of grandparents shared between cousins. For first cousins for example, *ϕ* is always 1/4 even if two, three, or four grandparents are shared by cousins; the exception being half first cousins where *ϕ* is 1/8. For more distant cousins, however, *ϕ* increases in multiple groups of four grandparents shared. For example, in second cousins, for two to four great-grandparents shared, *ϕ* is 1/16 (at 1/32 for half second cousins), while for five to eight great-grandparents shared it is 1/8. It proceeds similarly for increasingly distant cousins.

For (non-removed) cousins that share common ancestors *G* = *G*_*XA*_ = *G*_*Y A*_ generations in the past, *ϕ* can be calculated as

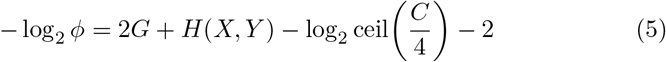

The function ceil in equation 5 is the ceiling function which rounds up to the smallest integer greater than or equal to *C/*4. Once we have *ϕ*, we can calculate the dominance correlation between cousins, 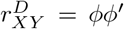 using *ϕ*′ = 2*r*_*XY*_ − *ϕ*. Table 2 shows how various degrees of relations can effect the additive and dominant correlations between different first through third cousins.

**Table 2.**
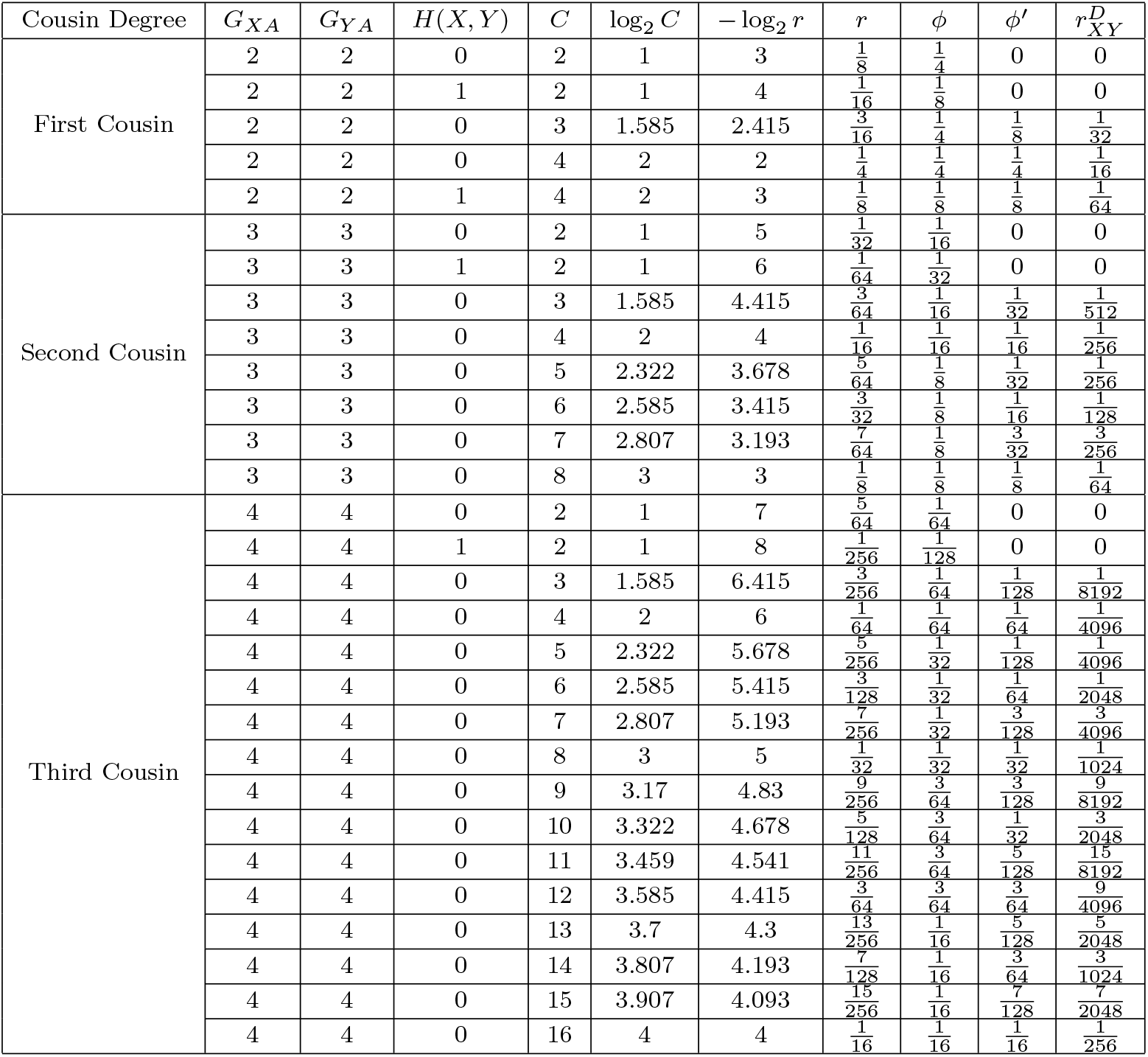
The additive and dominance correlations between various first, second, and third cousins using equations 2 and 4 where *C* is the number of shared relatives in the generation of the most recent common ancestor. Of note are double cousins where *G*_*X*_ = *G*_*Y*_ = 2 and *C* = 4 which share the same four grandparents and double half cousins which share the same variables except *H*(*X,Y*) = 1

## 3 Epistatic Covariances

The decomposition of the covariances of relatives who have epistatic genetic effects amongst loci in linkage equilibrium was given by [Kempthorne 1954, Kempthorne 1955, Cockerham 1954]. In short, for additive epistatic effects of two (AA), three (AAA), or n (*A*^*n*^) loci, the covariance is given by

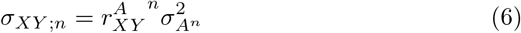

This can be represented in its effects by

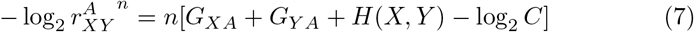

where *n* = 1 is the base case with no epistasis considered. In full siblings or cousins with dominance correlation, for *n* loci in linkage equilibrium that have *m* loci with additive epistasis and *n − m* loci with dominance epistasis the epistatic covariance for *X* and *Y* is

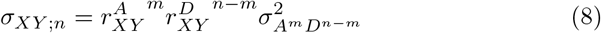

## 4 Results and Discussion

This paper does not present any new results regarding the kinship coefficients or correlations between relatives themselves. This is already a path well worn and verified by both simulation and increasingly, genomic data of large pedigrees. However, it does demonstrate that the correlations between arbitrary relatives can be quickly and easily calculated in a non-recursive fashion taking into account a few aspects of the pedigree tree for the relatives in question. While with modern computers, computational overhead is not much of an issue, and tables of kinship coefficients abound, this method may provide a shorthand for both researchers and students of quantitative genetics to ease the process of pedigree analysis.

